# Shifts in Genetic Diversity of Porcine Reproductive and Respiratory Syndrome Virus 2 in Vietnam Before and After African Swine Fever: Increased Diversity and Novel Sub-lineages

**DOI:** 10.64898/2026.07.06.736905

**Authors:** Thanh Che Nguyen, Nakarin Pamornchainavakul, João P. Herrera da Silva, Roongroje Thanawongnuwech, Kimberly VanderWaal

## Abstract

Porcine reproductive and respiratory syndrome virus 2 (PRRSV-2) remains one of the most important transboundary pathogens affecting swine production in Vietnam; however, it remains poorly understood how long-term evolutionary dynamics were impacted by the African swine fever (ASF) epidemic, a period of time where swine population demographics and movement were heavily perturbed. We investigated the molecular epidemiology, evolutionary history, and phylogeographic dynamics of PRRSV-2 circulating in Vietnam between 2007 and 2024 by integrating 366 Vietnamese ORF5 sequences with a globally curated lineage reference. Maximum-likelihood phylogenetic, Bayesian phylodynamic, and discrete phylogeographic analyses revealed that the Vietnamese PRRSV-2 population underwent substantial reshaping after the ASF epidemic, shifting from a predominantly endemic sub-lineage L8E population to a genetically diverse viral community comprising multiple established and newly emerging sub-lineages. Despite these epidemiological changes, the endemic sub-lineage L8E population maintained a relatively stable evolutionary rate across the pre- and post-ASF periods, suggesting that ASF reshaped viral population structure rather than intrinsic evolutionary dynamics. Two previously unclassified viral clusters circulating in Vietnam and Thailand fulfilled all criteria for formal designation and were recognized as the novel sub-lineages L1M and L10B by the International PRRSV-2 Nomenclature Consortium. Phylogeographic reconstruction further demonstrated contrasting transmission patterns among major sub-lineages, including long-term endemic persistence of L8E, repeated unidirectional introductions of sub-lineages L1M and L10B from Thailand, and bidirectional transpacific dissemination of sub-lineage L1A linking Southeast Asia and North America. Collectively, these findings demonstrate that the ASF epidemic coincided with a fundamental reshaping of the PRRSV-2 epidemiological landscape in Vietnam while revealing Southeast Asia as an active center of ongoing viral diversification. This study provides an updated evolutionary framework for PRRSV-2 surveillance and highlights the importance of continuous genomic monitoring and regional collaboration for the early detection and control of emerging transboundary variants.

## 1. Introduction

Porcine reproductive and respiratory syndrome (PRRS) remains a primary threat to sustainable swine production across Southeast Asia, inflicting severe reproductive failure in sows and respiratory distress in growing pigs [1]. The etiological agent, porcine reproductive and respiratory syndrome virus (PRRSV), is an enveloped, single-stranded positive-sense RNA virus belonging to the family *Arteriviridae* (order *Nidovirales*) [2]. Its genome length is roughly 15 kilobases, contains at least ten open reading frames (ORFs): ORF1a, 1b, 2a, 2b, and 3 to 7, encoding eight structural proteins: glycoproteins (GP2-5 and GP5a), envelope (E), matrix (M), and nucleocapsid (N) [3]. The lack of proofreading activity in its viral RNA-dependent RNA polymerase and frequent inter- and intra-lineage recombination events drive the continuous emergence of novel immunologically evasive variants and complicate control strategies [4, 5]. The virus is classified into two distinct species: *Betaarterivirus europensis* (PRRSV-1, historically European genotype) and *Betaarterivirus americense* (PRRSV-2, historically North American genotype) [2]. PRRSV-2 has been the dominant virus type in Asia, particularly in Vietnam, since the first emergence of the highly pathogenic (HP-PRRSV) lineage L8 in 2007 [6].

To systematically track this genetic hyper-variability, the global scientific community relies heavily on genetic characterization of the ORF5 gene, which encodes the major viral glycoprotein 5 (GP5) [7, 8]. PRRSV-2 ORF5 exhibits extraordinarily high substitution rates (10^-3^ to 10^-2^ substitutions/site/year, s/s/y) [7, 9]. A newly standardized global nomenclature system, curated by the International PRRSV-2 Nomenclature Consortium (IPNC, https://github.com/PRRSV-2-Nomenclature-Consortium/International-PRRSV-2-Nomenclature-Consortium/), utilizes explicit molecular criteria optimized from previous studies to classify PRRSV-2 into distinct phylogenetic lineages and sub-lineages based on pairwise nucleotide distances and phylogenetic analysis [7, 8, 10–12]. Through the IPNC framework, novel lineages and sub-lineages have been reviewed and endorsed, including lineage L12 in South Korea [13] and sub-lineages L1K and L1L in North America [14]. In sum, there are at least 12 lineages (L1-L12) and 23 sub-lineages for PRRSV-2 ORF5 sequences globally.

Vietnam is Southeast Asia’s leading pig producer, with 3.94 million metric tons in 2025 [15]. PRRSV-2 emerged in early 2007 and has been the dominant species causing long-term economic burdens associated with reproductive failure and reduced growth performance [16]. The epidemiological landscape of PRRSV-2 has undergone profound shifting over the past 20 years. However, there have been no comprehensive updates on the evolutionary dynamics of local strains since 2015, leaving nearly a decade of evolution and spatiotemporal dynamics uncharacterized. During 2007 - 2015, the predominant HP-PRRSV sub-lineage had an estimated evolutionary rate of 4.46 x 10^-3^ s/s/y, and the time of the Most Recent Common Ancestor (tMRCA) was in the early 2000s [17]. As a genetically variable and endemic virus, relying solely on historical sequences severely undercuts the ability to detect newly emerging variants or hidden transboundary introductions in real time.

This evolutionary landscape has potentially been influenced by the massive shift in the swine industry caused by the African swine fever (ASF) epidemic, which has fundamentally altered swine industry structures and generated specific selective pressures that have reframed PRRSV-2 evolution. The high mortality and subsequent culling during an ASF crisis trigger sharp declines in the live pig population and prompt immediate enhancements in farm biosecurity and shifts in animal movement patterns [18, 19]. This sudden contraction and strict containment could induce temporary viral genetic bottlenecks, dramatically shifting viral genetic pools. Indeed, post-ASF genomic surveillance of PRRSV-2 in China revealed that intensive biosecurity and animal transport restrictions appeared to suppress regional inter-lineage recombination while driving local intra-lineage recombination within isolated clusters, accelerating lineage displacement, and facilitating complex recombination events [20].

Therefore, in Vietnam, establishing an updated and rigorous molecular surveillance framework is urgently needed to address critical gaps in current epidemiological data, characterize the structural shifts induced by epidemic field pressures, and standardize lineage classification within an increasingly complex viral ecosystem. This study aims to perform an extensive epidemiological and phylogenetic analysis of PRRSV-2 strains circulating in Vietnam from 2007 to 2024. By leveraging contemporary genomic data, this study could map the shifting patterns of (sub-)lineage dominance, propose new sub-lineages in the post-ASF swine landscape, and provide an updated, standardized baseline essential for national disease control and management strategies.

## 2. Materials and Methods

### 2.1. Compilation of Global and Domestic PRRSV-2 ORF5 Sequence Data

The initial dataset contained 32,679 PRRSV-2 ORF5 sequences, obtained from two principal repositories spanning a 24-year (1990-2024) window **(Supplementary Figure S1)**. First, a curated baseline of 7,905 reference sequences was assembled and classified according to recognized modern lineage nomenclature guidelines [10]. This reference set integrated published sequences, encompassing 1,078 designated lineage anchor sequences described by Yim-im et al. [10], 690 representative North American sequences from Paploski et al. [8], and 6,137 Chinese sequences compiled by Tian et al. [21]. Second, a comprehensive global dataset comprising 24,774 sequences was extracted from the United States Swine Pathogen Database (US-SPD, https://swinepathogendb.org, accessed on February 02, 2026), utilizing strict search criteria: Organism: “Porcine reproductive and respiratory syndrome virus-2”, Sequence length: “603”, Annotations: “ORF5 full”, and “Omit duplicate sequences”.

Contemporary Vietnamese PRRSV-2 ORF5 records were retrieved from the National Center for Biotechnology Information (NCBI) GenBank database. Queries were conducted using combined search keywords “Porcine reproductive and respiratory syndrome virus 2” OR “PRRSV-2”, AND “Viet Nam”, AND “ORF5 gene” OR “GP5 protein”, indexed up to February 02, 2026. Consequently, 366 PRRSV-2 ORF5 sequences recovered from 2007 to 2024 were retained for preliminary analysis **(Supplementary Figure S1 and Figure 1**).

**Figure 1.**
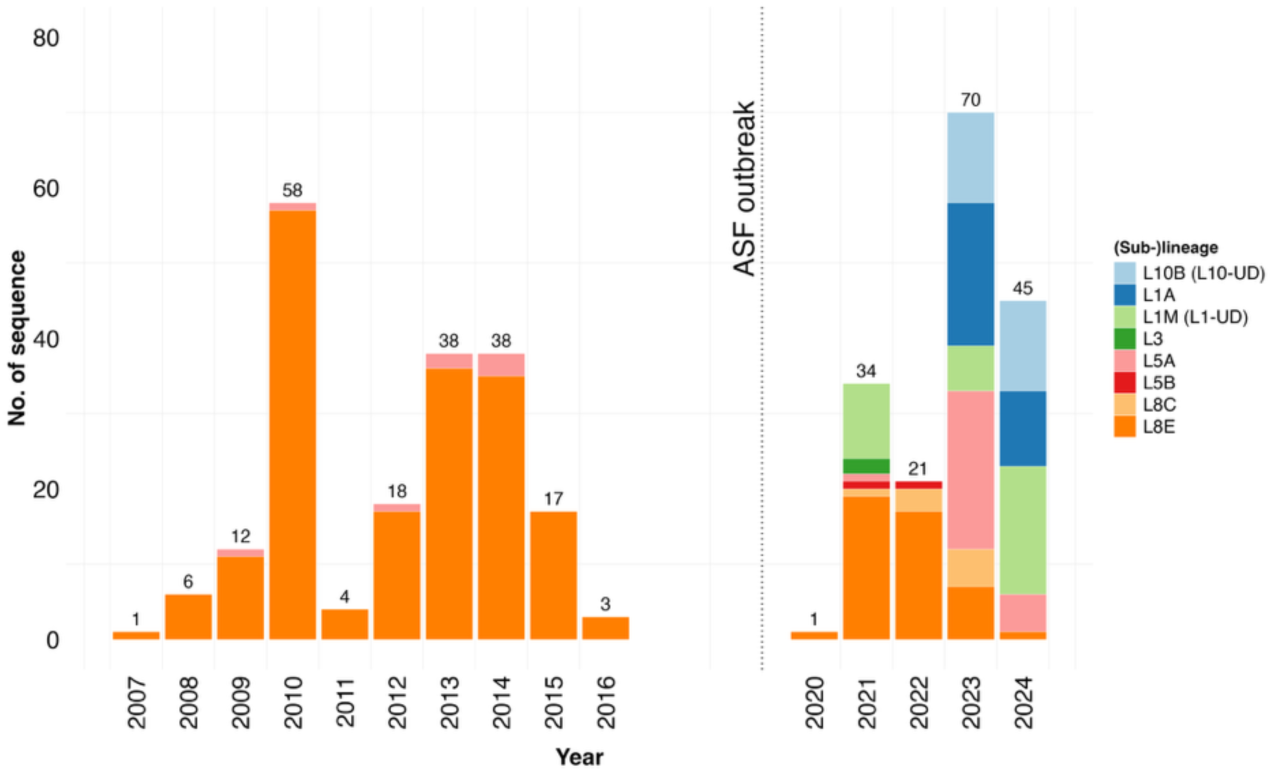
Numbers of PRRSV-2 ORF5 Vietnamese sequences collected across two distinct epidemiological windows: pre-ASF (2007 – 2016) and post-ASF (2020 – 2024) epidemic. Numbers above the columns display the absolute sequence counts per year. The dotted line represents the 2019 ASF outbreak in Vietnam. Colors indicate the phylogenetic sub-lineage, with “UD” denoting monophyletic clades that were classified as undetermined by PRRSLoom.

### 2.2. Sequence Quality Control and Filtering

The entire dataset, including global reference and Vietnamese sequences (*n* = 33,045), was aligned using Multiple Alignment using Fast Fourier Transform (MAFFT) v7.490 [22]. The nucleotide-aligned dataset was subjected to a rigorous quality-control and filtering pipeline to remove substandard sequences before downsampling. Entries lacking sample collection date and location (*n* = 10,842) were excluded. Duplicate sequences, defined by records sharing the same GenBank accession numbers or exhibiting 100% nucleotide identity with identical metadata (sampling time and location), were collapsed into a single representative sequence (*n* = 6,467). Sequences displaying unresolved nucleotide ambiguities (apart from A, T, G, and C) or deviating from the expected 603 bp alignment length (*n* = 3,390) were removed. Following initial filtering, sequences from European countries (*n* = 337) were excluded due to sparse data density. The final dataset (*n* = 12,009) comprised the Vietnamese sequences (*n* = 366) and global references (*n* = 11,643) originating from the USA, Canada, Mexico, South Korea, Japan, China, Taiwan, and Thailand (**Supplementary Figure S1)**. These retained sequences were utilized for downstream analysis, and their initial operational lineage was identified via the PRRSLoom analytical interface (https://stemma.shinyapps.io/PRRSLoom/) [11]. Vietnamese sequences were classified as pre- versus post-ASF outbreak using year of sampling relative to the date of ASF introduction (2019) (**Figure 1**).

### 2.3. Global Reference Dataset Downsampling

To establish a computationally tractable and epidemiologically balanced dataset for molecular epidemiological analysis, a stratified downsampling workflow adapted from Pamornchaivanavakul et al. [9] was applied to the 11,643 reference sequences. This stratified downsampling approach mitigated spatial (geographic locale) and temporal (time of collection) sampling biases that can distort phylogeographic analysis, while preserving the underlying global diversity. Contemporary wild-type sequences originating in Vietnam and countries with fewer than 50 total records were exempted from downsampling to maintain localized genetic resolution. Downsampling was performed three times with replacement, generating three independent sub-datasets with possible sequence overlap, which were analyzed separately downstream. Details of downsampling are described below.

#### 2.3.1. Subsampling for Phylogenetic Lineage Assignment and Recombination Detection

The global reference dataset was partitioned into distinct spatiotemporal strata based on country and collection year. Within each independent country-year stratum, pairwise genetic similarity matrices were computed across full alignments using the ‘ape’ package in R [23]. Sequences sharing a genetic similarity threshold of ≥ 98% within a given stratum were pooled into discrete micro-clusters, and a single representative sequence from each cluster was preserved [24, 25]. When multiple sequences had identical, fully complete metadata records (i.e., same country and year), the algorithm selected the centroid sequence of the micro-cluster as the representative sequence. Following this triple-iterated subsampling step, the resulting sub-datasets were utilized to construct maximum-likelihood (ML) phylogenetic trees (described in Section 2.4) to refine the classification and identify potential recombination events (described in Section 2.5).

#### 2.3.2. Subsampling for L1A-specific Phylogeographic Reconstruction

Preliminary analysis revealed a cluster of sub-lineage L1A sequences that occurred across Vietnam and the USA, which was surprising and warranted closer investigation. To trace potential transboundary migration routes and ancestral spatial origins of sub-lineages L1A in Vietnam, we constructed a L1A-specific dataset for phylogeographic analysis. To account for imbalances in data availability across countries, a weighted random sampling algorithm was used in place of standard uniform sampling methods. Given the limited number of Vietnamese sub-lineage L1A sequences (< 30), sequences from other countries were capped at twice the Vietnamese sample size to maintain geographic balance, except for Taiwan, where only 12 sequences were available. Sampling weights were then assigned to each individual sequence *(i)* within a defined spatiotemporal stratum. The sampling probability weight *(W_i_)* was calculated inversely proportional to its raw historical metadata density, as follows:

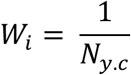

where *N_y.c_* represents the absolute sequence volume within a given collection year *(y)* and country *(c)*. By equalizing selection probabilities across periods of sparse and dense data collection, this sampling scheme downweighed heavily sequenced epidemic peaks to prevent temporal sampling bias [26].

This subsampling was replicated three times and analyzed separately to ensure robustness of results across independent sub-samples. The final sub-dataset comprises up to 225 sequences from five nations (USA, South Korea, China, Taiwan, and Vietnam) for discrete trait phylogeographic modeling.

### 2.4. Maximum-Likelihood Phylogenetic Analysis and Genetic Distances Estimation

The final down-sampled datasets (i.e., global subsampling dataset [*n* = 2,264] and all 366 Vietnamese sequences) were aligned as mentioned in Section 2.2. ML trees were inferred using IQ-TREE v2.4.0 [27] with 1,000 ultrafast bootstrap replicates to evaluate clade support. A best-fit GTR + I + R10 nucleotide substitution model was selected via ModelFinder, based on the minimum scoring thresholds of Bayesian Information Criterion [28]. Crucially, many sequences initially categorized as “undetermined” by the automated PRRSLoom classifier were observed to fall within two main monophyletic clades [8, 10]. Bootstrap values of the ancestral nodes of each clade were evaluated. Phylogenetic trees and taxonomic annotations were visualized and mapped using the ‘ggtree’ package in R [29].

To estimate genetic divergence between the two undetermined Vietnamese clades and the global reference sequences, a pairwise genetic distance matrix was calculated using the p-distance model in the Sequence Demarcation Tool v1.2 [30], based on a muscle alignment. The global reference dataset compiled by Herrera da Silva et al. [14] was used for this comparison, as this dataset included the most recently established L1 sub-lineages. Pairwise genetic distance profiles were plotted using the ‘ggplot2’ package in R software [31].

### 2.5. Screening for Recombination

Alignments incorporating the downsampled reference sets and the Vietnamese sequences were scanned to identify signals of intragenic and inter-lineage recombination using the Recombination Detection Program (RDP) v5.16 [32]. Seven exploratory algorithms were executed simultaneously: RDP [33], GENECONV [34], Chimaera [35], MaxChi [36], BootScan [37], SiScan [38], and 3Seq [39]. A potential recombination event was considered when statistical support was confirmed with a *p*-value < 0.05 across at least four independent methods. To prevent distortion of molecular clock rates and topologies, recombinant sequences were excluded from the dataset prior to Bayesian molecular-clock inference (described in 2.7) and ancestral-state modeling (described in 2.8).

### 2.6. Sub-lineage-specific Analysis and Rationale

To investigate the spatial epidemiology and evolutionary trends of wild-type strains, PRRSV-2 vaccine-like sequences were excluded from the dataset (*n* = 81 of 2,264) and the Vietnamese dataset (*n* = 42 of 366). Sequences demonstrating a pairwise nucleotide identity (described in Section 2.4) threshold exceeding 95% relative to the vaccine reference sequences were considered vaccine-like [10]. Six major global commercial vaccines were considered: Prevacent^®^ from Elanco US Inc. PRRS (L1D, KU131568), PRRSGard^®^ from Pharmagate Animal Health (L1F, OR293983), Ingelvac PRRS^®^ MLV from Boehringer Ingelheim Animal Health USA Inc. (L5A, AF066183), Prime Pac^®^ PRRS RR from Merck Animal Health (L7, AF066384), Ingelvac PRRS^®^ ATP from Boehringer Ingelheim Animal Health USA Inc. (L8A, AY656991), and Fostera^®^ PRRS from Zoetis Inc. (L8C, LC557039). Following vaccine exclusion, a finalized sub-dataset of 2,183 reference sequences and 324 Vietnamese sequences was retained for phylodynamic analysis.

Phylodynamic and phylogeographic modeling was restricted to the four major sub-lineages circulating in Vietnam (L8E, L1A, and two novel sub-lineages), while minor co-circulating clades with sparse longitudinal data were not analyzed (**Supplementary Figure S1**). The analytical frameworks were tailored to fit each group’s unique epidemiological context: we implemented a temporal Bayesian framework for sub-lineage L8E to monitor long-term domestic evolution across pre- and post-ASF era (section 2.7); performed phylodynamic and phylogeographic analysis for sub-lineage L1A to isolate its domestic tMRCA and map regional transboundary migration corridors (sections 2.7 and 2.8); and executed ancestral-tracking analysis of two novel sub-lineages by augmenting the Vietnamese datasets with their closest related sub-lineages to pinpoint the estimated timing and ancestral sources of their introduction (section 2.8).

### 2.7. Temporal Signal Evaluation and Bayesian Molecular Clock Inference

Prior to initiating Bayesian Markov Chain Monte Carlo (MCMC) simulations, the strength of the temporal evolutionary signal within each sub-lineage alignment was evaluated via root-to-tip regression analysis of the initial IQ-TREE topologies using TempEst v.1.5.3 [40]. Outlier sequences were removed using Tukey’s interquartile range (IQR) method [41]; Sequences with root-to-tip residuals exceeding 1.5 x IQR from the first or third quartiles were removed.

Bayesian ancestral inferences were performed in the Bayesian Evolutionary Analysis Sampling Trees (BEAST) v1.10.4 to model tMRCA parameters, ancestral population changes, and evolutionary rates [42]. The MCMC configuration utilized (i) the SRD06 codon-partitioned substitution model to account for varying evolutionary constraints across the protein-coding region by separating nucleotide positions 1+2 from position 3 [43]; (ii) an uncorrelated relaxed clock following a lognormal distribution [44]; and (iii) a flexible Gaussian Markov Random Field (GMRF) Bayesian SkyGrid coalescent tree prior to capture population size fluctuations [45]. MCMC chains were independently run three times for each sub-dataset for 300 million generations, sampling every 30,000 steps. Posterior log distributions from independent runs were inspected in Tracer v1.7.2 [46] to confirm convergence and combined using LogCombiner v1.10.4 [47]. Convergence required an Effective Sample Size (ESS) exceeding 200 for all primary parameters. Maximum Clade Credibility (MCC) trees were generated using TreeAnnotator v1.10.4 [47], discarding the initial 10% as burn-in. Finalized MCC trees were rendered in FigTree v1.4.4 [48] to display node heights, divergence dates, and posterior probabilities.

### 2.8. Discrete-space Ancestral State Reconstruction

To trace transboundary migration networks of sub-lineages L1A and two newly designated sub-lineages into and within Vietnam, discrete-space Bayesian phylogeographic models were implemented in BEAST v1.10.4 [42] using the sampling countries as discrete traits under an asymmetric continuous-time Markov chain (CTMC) matrix [49]. A Bayesian Stochastic Search Variable Selection (BSSVS) extension was applied to identify statistically significant transmission links. MCMC configurations, convergence criteria (ESS > 200), and tree summarization matched the parameters described in section 2.7. Directional transmission pathways were quantified via Bayes Factors (BF), and relative transition rates were calculated in SpreaD3 v0.9.7 [50]. To account for stochastic variation across independent runs, the final spatial trajectory for sub-lineage L1A was determined by calculating the arithmetic mean of the median transition rates across three independent sub-datasets.

## 3. Results

### 3.1. Dataset Curation, Sub-sampling, and Initial Lineage Classification

The final global dataset compiled for macro-level PRRSV-2 ORF5 classification was 2,630 sequences from 1991 to 2024. This dataset incorporated sequences from Canada (*n* = 287), China (*n* = 945), Japan (*n* = 22), Mexico (*n* = 71), South Korea (*n* = 207), Taiwan (*n* = 105), Thailand (*n* = 43), the USA (*n* = 579), Vietnam (*n* = 366), and vaccine references (*n* = 5). The original Vietnamese dataset (*n* = 366; collected between 2007 and 2024) was stratified into two distinct epidemiological periods relative to the introduction of ASF: pre-ASF outbreak phase (2007 – 2016, *n =* 195) and post-ASF outbreak phase (2020 – 2024, *n* = 171) to evaluate the impact of the ASF epidemic on PRRSV-2 evolutionary dynamics **(Figure 1)**. RDP analysis yielded no statistically significant evidence of intra- or inter-lineage recombinant events within the Vietnamese ORF5 sequences.

Classification using PRRSLoom initially partitioned the Vietnamese dataset into four established lineages (L1, L3, L5, and L8), and five sub-lineages (L1A, L1H-unclassified, L5A, L8C, and L8E). Intriguingly, 21 sequences were classified as sub-lineage L1H, a sub-lineage thought to be restricted to North American swine populations; however, an examination of these Vietnamese sequences relative to other L1H sequences in the phylogeny suggests that the PRRSLoom classification was erroneous. In addition, PRRSLoom classified 38 of the 366 sequences (10.4%) as “undetermined,” indicating that these sequences were not well-matched to any of the established sub-lineages.

### 3.2. ML Phylogenetic Topology and the Formalization of Novel Sub-lineages

To resolve the taxonomic status of the suspicious L1H-unclassified (*n* = 21) and undetermined (*n* = 38) sequences (hereafter referred to as the query sequences), global ML phylogenetic trees were constructed using IQ-TREE, incorporating anchor sequences for 12 recognized lineages (comprising 23 sub-lineages) identified by Yim-im et al. [10], Paploski et al. [8], and Herrera da Silva et al. [14]. The majority of established sub-lineages resolved as distinct monophyletic clades with robust bootstrap support (≥ 80), validating the evolutionary fidelity of the reference framework **(Figure 2)**. The 59 query sequences split cleanly from global (sub-)lineages. In particular, the “undetermined” sequences were partitioned into three distinct well-supported monophyletic clades (bootstrap values > 95): “Undetermined 1” (n = 2; sampled in 2021 and 2022) nested within the Chinese sub-lineage L5B (bootstrap = 96). “Undetermined 2” (n = 24; sampled in 2023 and 2024) diverged from a unique node basal to the lineage L10 references in Thailand (bootstrap = 100); this group was provisionally labeled “L10-UD”, and “Undetermined 3” (n = 12) combined with the 21 suspicious L1H-unclassified sequences from 2021 and 2023 – 2024 formed a well-supported clade nested within lineage L1 (bootstrap = 99), exhibiting a sister-group relationship with sub-lineage L1I from the USA; this group was provisionally labeled “L1-UD.” This group was not closely related to sub-lineage L1H **(Figure 2)**.

**Figure 2.**
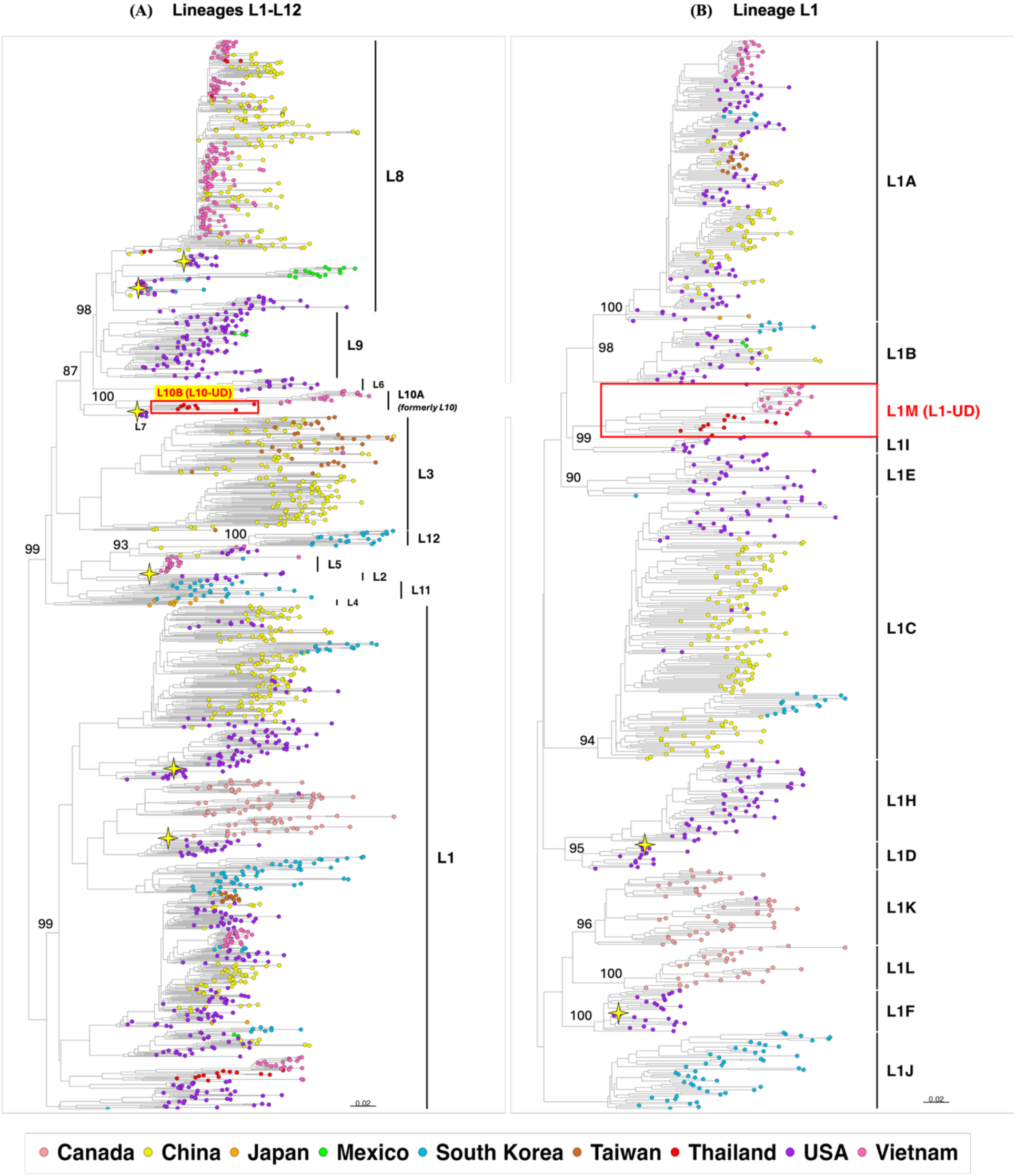
The maximum-likelihood (ML) phylogenetic tree analyses of PRRSV-2 Vietnamese sequences among **(A)** all 12 lineages (L1 – L12) and **(B)** 12 sub-lineages (L1A – L1J) within lineage L1. The yellowish 4-point stars present the vaccine references, and red labels and boxes indicate the proposed new sub-lineages. Bootstrap values above 80 are displayed at the major ancestral nodes.

We calculated pairwise nucleotide genetic distances between L1-UD and L10-UD against the comprehensive global reference set using SDT. Both candidate clusters exceeded the >10% genetic divergence typically observed between PRRSV-2 sub-lineage [10] (**Figure 2 and Supplementary Table S1**). The clades were reviewed by the IPNC (https://github.com/PRRSV-2-Nomenclature-Consortium/International-PRRSV-2-Nomenclature-Consortium/) and were endorsed as sub-lineage L1M and L10B. To maintain a systematic, hierarchical nomenclature, the previously un-subdivided ancestral lineage L10 sequences were reassigned as sub-lineage L10A.

Sub-lineage L10B exhibited a median intra-group genetic distance of 6.9% (IQR: 4.0 – 10.0%), reflecting substantial diversity. Its closest phylogenetic neighbor was lineage L10 (now sub-lineage L10A) (median distance: 12.8%, IQR: 12.4 – 13.1%) and sub-lineage L8C (13.9%, IQR: 13.4 – 14.6%), with distances to all other (sub-)lineages exceeded 14% **(Figure 3A and Supplementary Table S1)**. Sub-lineage L1M displayed a median intra-group genetic distance of 10.7% (IQR: 7.3 – 12.2%). Inter-group analyses showed that it is most closely related to sub-lineage L1I (median distance 13.9%, IQR: 13.1 – 14.4%), with all other comparisons exceeding 14% divergence **(Figure 3B and Supplementary Table S1)**. The genetic distance separating L10B and L1M was 18.4% (IQR: 17.2 – 19.1%), confirming their independent evolutionary trajectories.

**Figure 3.**
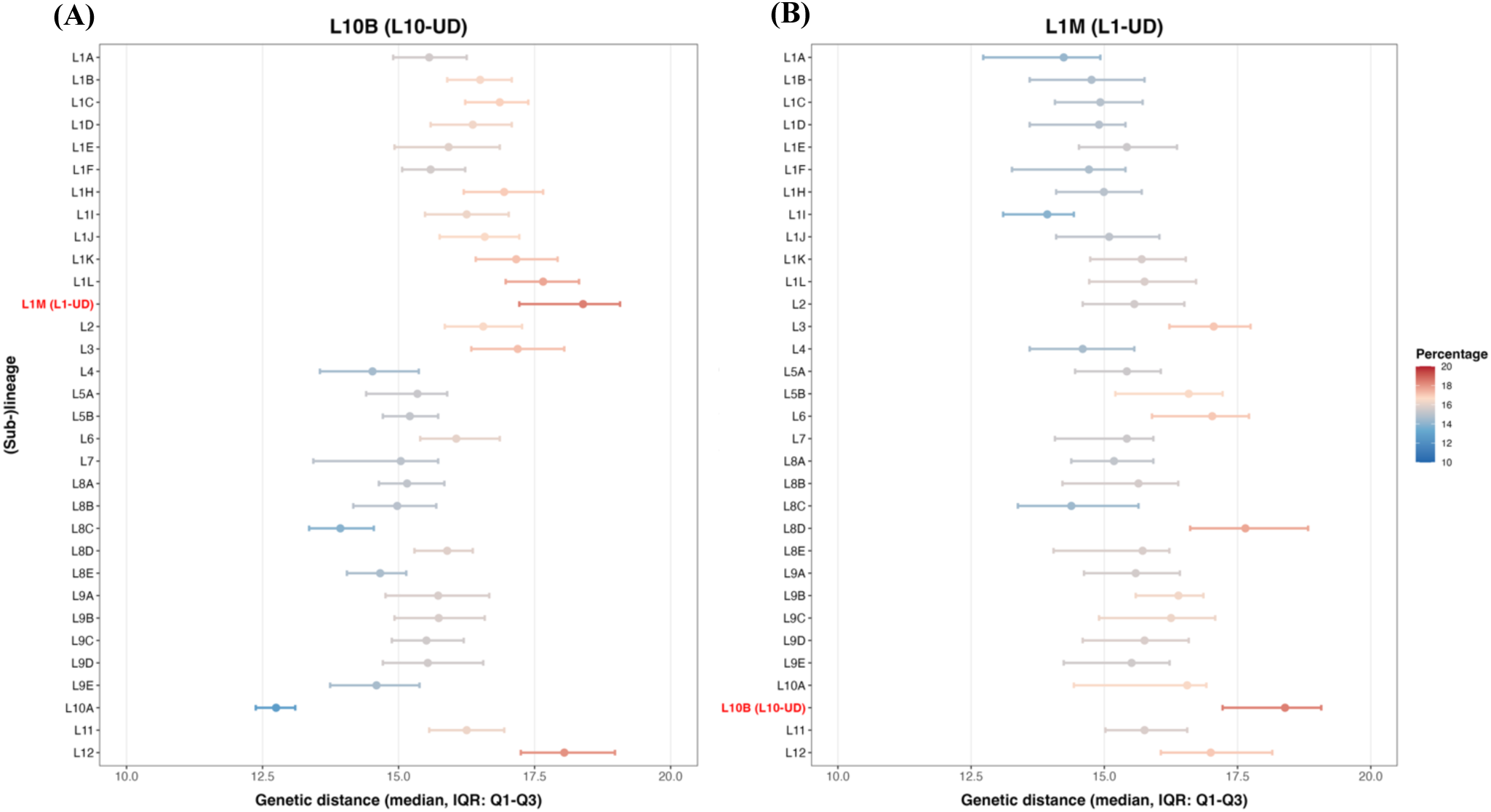
Comparative pairwise genetic distance analysis of novel PRRSV-2 sub-lineages: **(A)** L10B (L10-UD) and **(B)** L1M (L1-UD) relative to globally recognized PRRSV-2 (sub-)lineages. Dots and bars denote the median and interquartile range (IQR: Q1 – Q3), respectively. Nucleotide identities were calculated using SDT v1.2.

Spatial-temporal distribution data supported that these novel groups have circulated either solely in Vietnam or, more likely, in neighboring Thailand. While sub-lineage L10B was first described in Vietnam (2023 – 2024), its presence was recently corroborated by preliminary genomic surveillance data from Thailand in 2023 [51]. Similarly, the sub-lineage L1M was detected in Vietnam in 2021 and 2023 – 2024, but historical sequences demonstrate circulation in Thailand between 2008 and 2015 [51]. These parallel regional detections confirmed that these groups represent widespread evolutionary trends in Southeast Asia rather than localized or isolated sequencing artifacts.

### 3.3. Lineage Diversity Before and After African Swine Fever

Prior to ASF, our analysis demonstrates that sub-lineage L8E was predominant, accounting for 95.9% (187/195) of the pre-ASF dataset; the remaining 4.1% (8/195) was sub-lineage L5A and was likely associated with the Boehringer Ingelheim MLV (**Figure 1**). An expansion of lineage diversity was observed after ASF. Sub-lineages L8E (26.3%; 45/171) and L5A (15.8%; 27/171, of which 96.3% [26/27] were vaccine-like sequences) continued to circulate alongside a number of sub-lineages previously undetected in Vietnam. These include sub-lineages L1A (17.0%, 29/171), L8C (5.3%, 9/171, all of which were vaccine-like), L5B (1.2%, 2/171), L3 (1.2%, 2/171), and the newly established sub-lineages L1M (19.3%, 33/171) and L10B (14.0%, 24/171). Macro-level temporal analysis showed a wave of these introductions in 2021 for (sub-)lineages L1M, L5B, L8C, and L3, followed by the expansion of L1A and L10B in 2023 – 2024 (**Figures 1 and 2)**.

### 3.4. Bayesian Phylodynamic Modeling and Spatiotemporal Dynamics in Vietnam

To unravel the evolutionary history and introduction dynamics of the Vietnamese PRRSV-2 sub-lineages, molecular clock and discrete-trait phylogeographic analyses were performed using BEAST. Prior to analysis, 42 vaccine-like sequences sharing > 95% nucleotide identity with L5A (*n* = 34) and L8C (*n* = 9) commercial vaccines, representing 11.5% of the Vietnamese dataset, were removed to eliminate artificial evolutionary signals.

#### 3.4.1. Long-term Endemic Persistence and Evolutionary Rate of Sub-lineage L8E

As the only sub-lineage maintaining continuous endemicity across both investigated eras, sub-lineage L8E was evaluated to assess its evolutionary dynamics relative to the 2019 ASF epidemic. Under an uncorrelated relaxed lognormal molecular clock model, the historical baseline (pre-ASF era) mean nucleotide substitution rate was estimated at 2.04 x 10^-3^ s/s/y (95% highest posterior density [HPD]: 1.52 – 2.59 x 10^-3^). Analysis of the full longitudinal dataset (spanning both pre- and post-ASF eras) yielded a comparable mean substitution rate of 2.46 x 10^-3^ s/s/y (95% HPD: 1.95 – 3.00 x 10^-3^). The extensive overlap between these 95% HPD intervals indicated that the baseline evolutionary velocity of sub-lineage L8E remained highly stable in Vietnam, demonstrating that the severe macro-environmental disruptions and swine population culling caused by the 2019 ASF epidemic did not significantly alter the long-term accumulation of substitutions within this endemic sub-lineage.

tMRCA estimated for the root node of the Vietnamese L8E population was highly congruent between the pre-ASF dataset (estimated: 2003.9; 95% HPD: 1999.4 – 2006.6) and the full dataset (estimated: 2003.4, 95% HPD: 1996.3 – 2006.4) **(Supplementary Figure S2)**. This temporal consistency was topologically supported by the direct descendant relationships within the tree; rather than forming distinct, independent branches from the root, the post-ASF L8E field strains nested descendants within the clades that existed during the pre-ASF era. This continuous topological branching suggests unbroken domestic persistence and diversification of the endemic L8E population across the ASF epidemic threshold without any apparent population bottlenecks. However, given the restricted post-ASF L8E sequence availability, the potential for low-level, undetected secondary transboundary introduction cannot be completely ruled out in this study.

#### 3.4.2. Trans-Pacific Migration of Sub-lineage L1A

To model the spatiotemporal history of sub-lineage L1A, a sub-dataset of 225 PRRSV-2 ORF5 sequences from five countries (China, the USA, and South Korea: *n* = 61-62 per country, Taiwan: *n* = 12, and Vietnam: *n* = 29) were analyzed. Discrete-trait phylogeographic reconstruction revealed a complex, bidirectional transmission history between the USA and East/Southeast Asia **(Figure 4)**. The primitive origin of the L1A clade was situated within the US swine population in the early 2000s. The virus subsequently evolved and was exported westward, resulting in separate transboundary introduction into Vietnam in the mid-to-late 2010s (tMRCA: 2014.6; 95% HPD: 2014.1 – 2015.4), South Korea (tMRCA: 2017.4, 95% HPD: 2015.8 – 2018.9), Taiwan (tMRCA: 2016.6, 95% HPD: 2015.5 – 2017.4) and at least two introductions from the USA into China (tMRCA: 2013.9, 95% HPD: 2013.4 – 2014.7 and tMRCA: 2014.2, 95% HPD: 2013.8 – 2014.8). Incomplete sampling of sequences in Asia could affect this finding.

**Figure 4.**
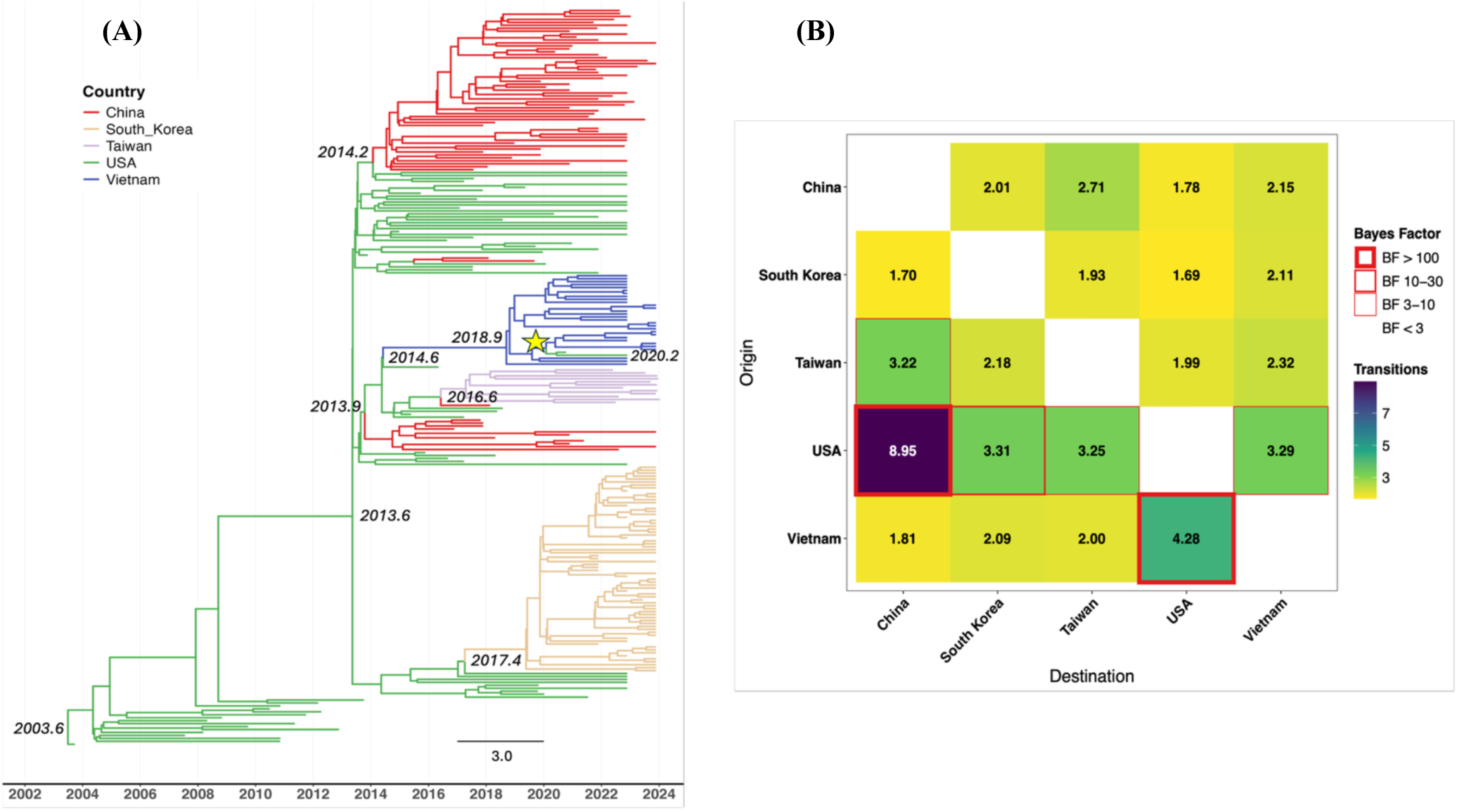
Spatiotemporal evolutionary dynamics of PRRSV-2 sub-lineage L1A. **(A)** Discrete-space diffusion Maximum Clade Credibility (MCC) phylogeographic tree reconstruction. Internal node labels represent the estimated time of the Most Recent Common Ancestor (tMRCA) for major clades. The yellowish star displays the reintroduction from Vietnam to the USA around 2020. **(B)** Matrix heatmap detailing transboundary transition counts. Red borders denote the magnitude of Bayes factor support; only well-supported transitions (BF ≥ 3) are outlined.

Following the introduction of the virus into Vietnam, it established stable endemic transmission, leading to a major internal diversification event around 2018 (2018.9, 95% HPD: 2017.8 – 2019.8) within the Vietnamese swine population. Crucially, the MCC tree captured an unexpected reverse-migration event back to North America: an evolutionary branch diverged from the domestic Vietnamese pool between 2019.5 and 2020.0, transitioning directly back into the USA swine population **(Figure 4A)**. This reintroduction coincided with the recent variant L1A.11 observed between 2020 and 2025 in the USA [11]. The earliest detection of L1A.11 in the USA (GenBank accession no. PV967014) was in 2020 and shared 97.8% nucleotide identity with the closest Vietnamese sub-lineage L1A sequence. Moreover, the representative L1A.11 sequence (GenBank accession no. OR293446) exhibited a median genetic distance of 2.3% (minimum 1.2%) relative to the Vietnamese sub-lineage L1A sequences. Consistent with these observations, all 29 sub-lineage L1A sequences were subsequently classified as variant L1A.11 using PRRSLoom-Variants (https://stemma.shinyapps.io/PRRSLoom-variants/) [11]. This date aligns with the inferred date of the Vietnam-U.S. transition from our analysis (2019.5 – 2020.0), but was beyond the scope of this study to further investigate this trans-Pacific migration event.

This transboundary spread between the USA and Southeast Asia was robustly supported by discrete-trait modeling, which identified asymmetric trans-Pacific migrations (BF > 100, strong statistical support). The primary historical expansion route was from the USA to China, with the highest viral inferred migration events (8.95 transitions). The USA was also inferred as the source of sub-lineage L1A for South Korea (3.31 transitions), Vietnam (3.29 transitions), and Taiwan (3.25 transitions; all BF 10–30, substantial support). Crucially, the model provided strong support for reverse-migration from Vietnam back to the USA (4.25 transitions, BF > 100) **(Figure 4B)**. This finding was consistent across three runs per sub-sample in three different sub-samples.

#### 3.4.3. Introduction of Sub-lineage L1M to Vietnam

The newly designated sub-lineage L1M followed a unidirectional dispersal pattern from Thailand to Vietnam **(Figure 5)**, though sampling availability could affect this finding. To better define the ancestral root of this sub-lineage, sequences from the closest related sub-lineage (L1I) were included in the analysis. The root of the L1M clade was localized to Thailand in the late 1990s (tMRCA: 1997.5; 95% HPD: 1992.5 – 2002.1), establishing a decade-long history of cryptic diversification prior to regional dissemination. While a Southeast Asia ancestral location seems likely for sub-lineage L1M, the exact country is difficult to determine due to uneven sampling across time and space.

**Figure 5.**
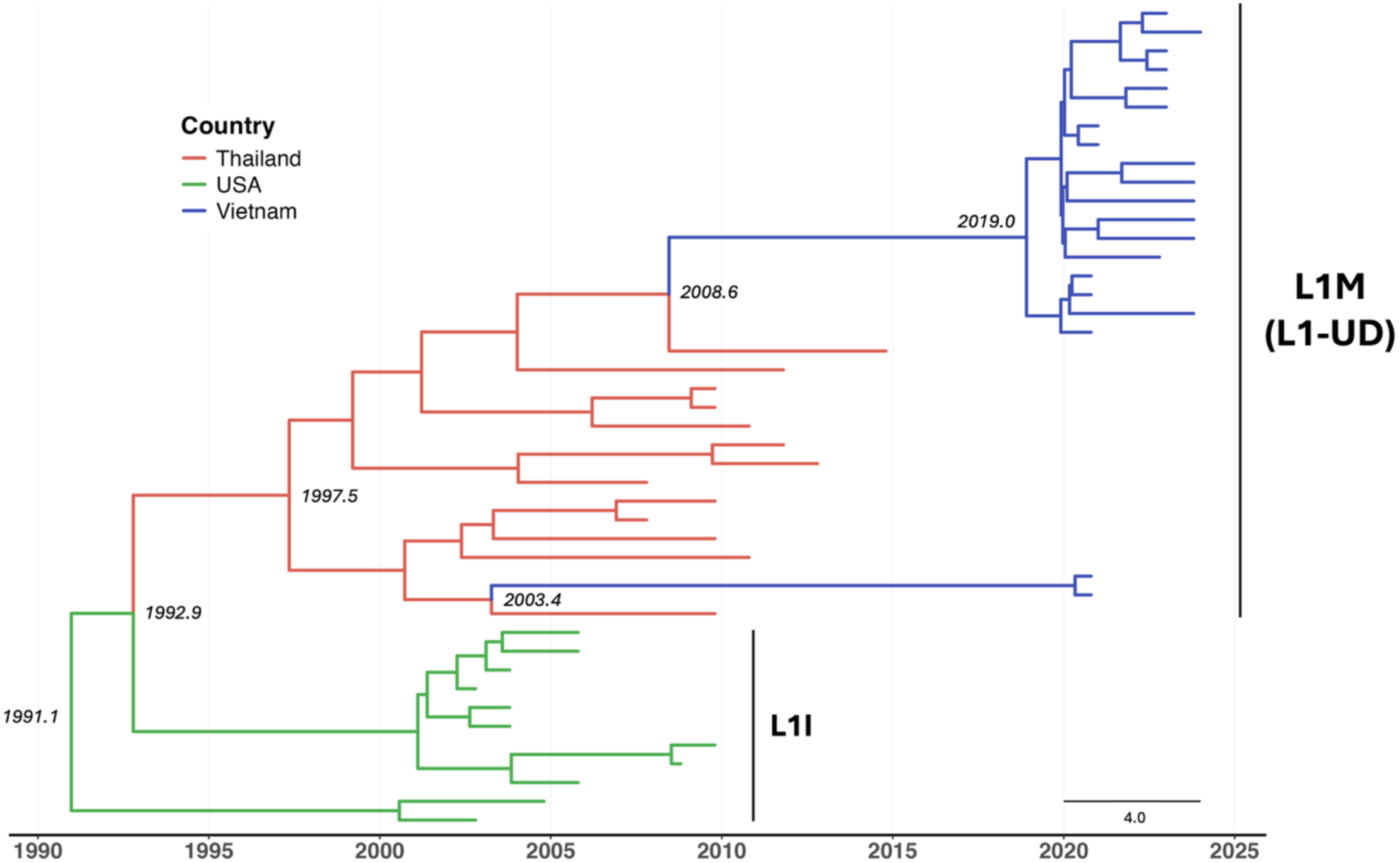
Maximum Clade Credibility (MCC) tree of PRRSV-2 Lineage L1 (L1M [L1-UD] and L1I). The tree was reconstructed using a Bayesian MCMC approach in BEAST v1.10.4. Branches are color-coded by geographic location as described in the legend. Internal node labels represent the estimated time of the Most Recent Common Ancestor (tMRCA) for major clades.

Rather than entering Vietnam via a single seeding event, the MCC tree suggests multiple seeding from Thailand to Vietnam, though lack of sequencing during the early 2010s makes it difficult to fully elucidate transboundary transmission dynamics. That being said, there appears to be a major introduction from Thailand to Vietnam sometime between 2008 and 2019. Given that sub-lineage L1M was not detected in Vietnam pre-ASF (before 2019), it seems likely that the introduction occurred closer to 2019.

#### 3.4.4. Introduction and Post-ASF Expansion of Sub-lineage L10B in Vietnam

The spatiotemporal dynamics of the newly designated sub-lineage L10B closely mirrored the apparent unidirectional, transboundary pattern observed in L1M, with a single seeding event from the Thai sub-lineage L10A population **(Figure 6)**. However, the lack of sequence data in the intervening years makes a definitive conclusion challenging. The global tMRCA of lineage L10 as a whole was estimated at approximately 2006 (tMRCA: 2005.8; 95% HPD: 2002.2 – 2008.4). Shortly thereafter, around 2008 (tMRCA: 2007.9; 95% HPD: 2005.5 – 2009.5), sub-lineage L10B began diverging.

**Figure 6.**
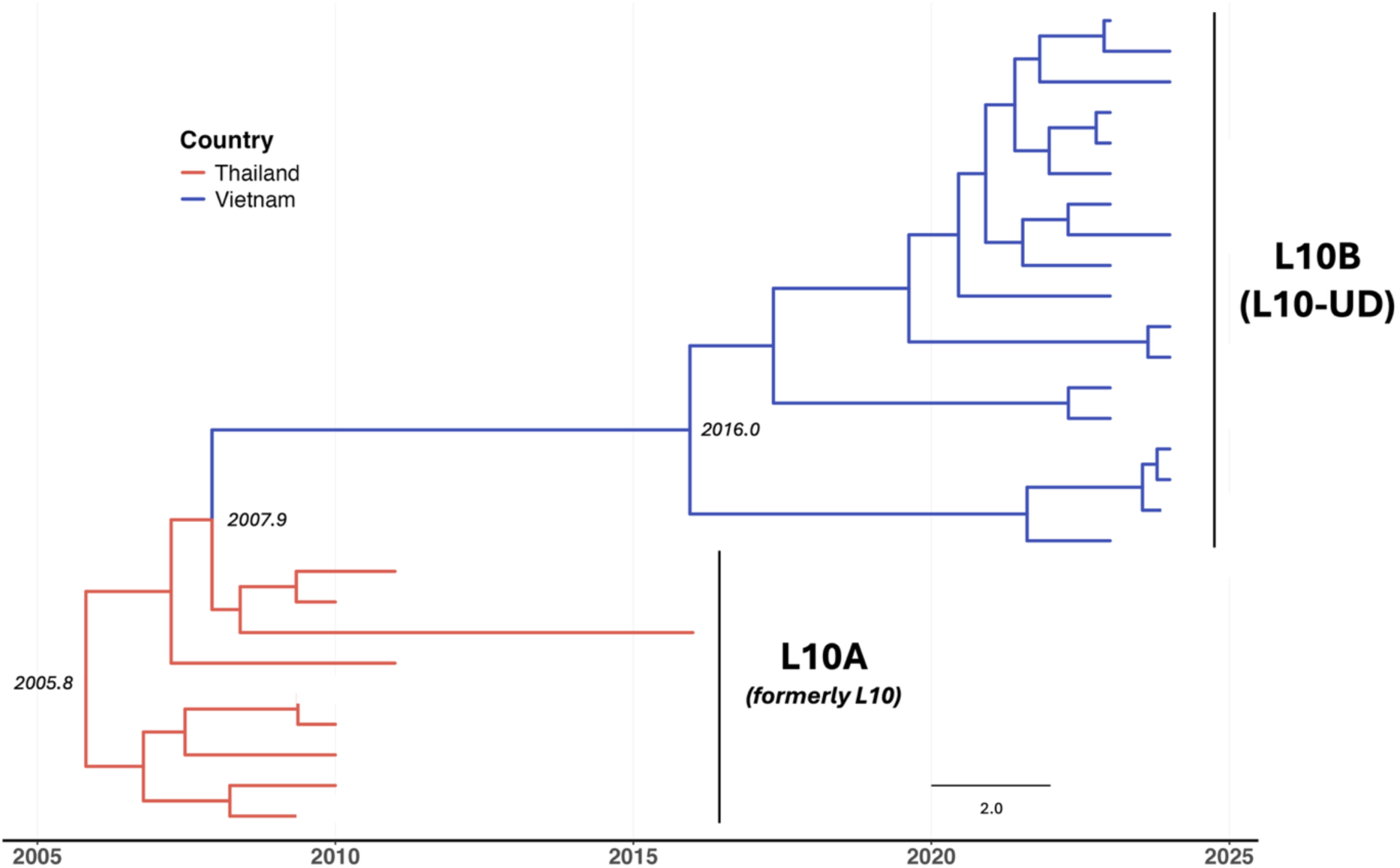
Maximum Clade Credibility (MCC) tree of PRRSV-2 Lineage 10 (L10A [formerly L10] and L10B [L10-UD]). The tree was reconstructed using a Bayesian MCMC approach in BEAST v.1.10.4. Branches are color-coded by geographic location as described in the legend. Internal node labels represent the estimated time of the Most Recent Common Ancestor (tMRCA) for major clades.

Based on available data, sub-lineage L10B spread from Thailand into Vietnam sometime between 2008 and 2016, though sub-lineage L10B did not appear in the Vietnamese pre-ASF dataset. Sub-lineage L10B underwent a major domestic expansion event beginning around 2016 (tMRCA: 2016.0; 95% HPD: 2011.4 – 2019.6) and possibly as late as July 2019, which would be compatible with a post-ASF expansion driven by animal movement to meet increased market demand **(Figure 6)**.

## 4. Discussion

PRRS is one of the most economically devastating and challenging endemic diseases in the global swine industry, including Vietnam [16]. PRRSV-2 continues to evolve rapidly in response to changing production systems, animal movement, and host immunity, making continuous molecular surveillance essential for disease control, especially after ASF incursion [8]. This study provides the first comprehensive characterization of lineage diversity and evolutionary dynamics of PRRSV-2 in Vietnam, spanning nearly two decades (2007 – 2024). Our results demonstrate that the Vietnamese PRRSV-2 population has undergone substantial reframing following the ASF epidemic, transitioning from a predominantly L8E-endemic population (2007 – 2016), as previously reported [6, 17, 52], to a more genetically heterogeneous viral community comprising multiple established and newly emerging sub-lineages (2020 – 2024). Importantly, this study formally identified two previously undescribed PRRSV-2 sub-lineages, L1M and L10B. Together, these findings establish an updated evolutionary baseline for PRRSV-2 in Vietnam.

One of the most notable findings was that the ASF epidemic appeared to reshape the epidemiological landscape of PRRSV-2 without substantially altering the evolutionary rate of the long-established endemic sub-lineage L8E. Although ASF caused unprecedented reductions in pig populations through mortality and large-scale culling, together with stricter biosecurity and movement restrictions, the estimated substitution rate of L8E remained consistent before and after the ASF epidemic. This observation suggests that the intrinsic molecular evolutionary rate of an established PRRSV-2 lineage is relatively resilient to short-term demographic perturbations. Instead, post-ASF L8E viruses appeared to evolve primarily through continued diversification of existing domestic clades rather than through repeated replacement by newly introduced variants. Similar observations have been reported in China, where ASF-associated movement restrictions reduced inter-provincial dissemination of NADC30-like (sub-lineage L1C) strains [20]. Collectively, these findings indicate that ASF did not directly accelerate PRRSV evolution; rather, it modified the ecological conditions under which different viral populations were able to persist, spread, and diversify.

The post-ASF period was characterized by the emergence and successful establishment of several previously uncommon or newly introduced PRRSV-2 sub-lineages. This increase in viral diversity likely reflects changes in pig production following ASF, including herd depopulation, rapid repopulation with replacement breeding stock, restructuring of production systems, and renewed animal movement [53]. By 2024, the total number of national pig heads recovered was approximately 27 million, and the number of smallholder farms dropped by almost 1 million compared with 2018, reflecting the consolidation of pig production into larger commercial systems, which were better able to recover from ASF [54]. However, biosecurity and quarantine measures vary across these production systems, potentially resulting in repeated viral introductions and establishment [19]. Although our phylogenetic results differed from those previously reported in China regarding genetic diversity, shifts in PRRSV-2 lineage composition were similar: the prevalence of sub-lineage L1A (NADC34-like) viruses increased markedly after ASF, while sub-lineage L8E (HP-PRRSV-like) strains became comparatively less dominant [20]. However, to be more confident, further analyses with a larger number of sequences, particularly in the post-ASF period, are warranted. Together, these observations suggest that major disease outbreaks can indirectly reshape the genetic diversity of other endemic pathogens by altering host population structure and transmission networks.

A major contribution of this study is the formal identification of two novel PRRSV-2 sub-lineages, L1M and L10B. These viruses fulfilled all criteria established by the IPNC, including robust monophyletic clustering, appropriate within- and between-(sub-)lineage genetic divergence, and sustained epidemiological circulation [13]. Their recognition further validates the robustness of the recently standardized global nomenclature framework, which improves reproducibility and consistency of PRRSV classification across studies [10–12]. Indeed, at the time of publication, the two new sub- lineages have been incorporated into PRRSLoom so that others will be able to easily assess whether their sequences belong to these new clades. Molecular clock analyses further suggested that both sub-lineages L1M and L10B have likely circulated cryptically for many years before being formally recognized. This discrepancy between estimated emergence and first detection most likely reflects historical limitations in genomic surveillance (i.e., the sparse availability of viral genomes in public sequence databases) rather than recent viral evolution. Similar retrospective identification of long-circulating novel PRRSV-2 sub-lineages has been reported in Canada (sub-lineages L1K and L1L) and South Korea (lineage L12) following expansion of molecular surveillance data [13, 14]. These findings emphasize that the current diversity of PRRSV-2 circulating throughout Southeast Asia is likely still underestimated, highlighting the importance of continuous genomic surveillance and regional data sharing.

The phylogeographic analyses further demonstrated that different PRRSV-2 sub-lineages follow distinct transmission dynamics, although the lack of consistently available sequence data across space and time may influence inferred spatiotemporal patterns [55]. Both sub-lineages L1M and L10B exhibited predominantly unidirectional introduction from Thailand into Vietnam, followed by successful domestic establishment and diversification. These findings support the hypothesis that cross-border livestock movements remain an important mechanism for regional PRRSV dissemination in mainland Southeast Asia. Although legal importation of breeding pigs is regulated, informal movement of live pigs, breeding materials, contaminated vehicles, and other indirect transmission pathways may facilitate repeated viral introduction across neighboring countries [56, 57]. To cool down pork prices driven by the first ASF wave in Vietnam, approximately 5 million Thai live pigs were imported from Thailand in 2020; such movements were banned a year later due to the ASF outbreak in Thailand [58]. The apparent absence of reverse transmission for these two sub-lineages suggests that Vietnam functioned primarily as a recipient rather than a source population for their regional spread, though results could differ with the inclusion of additional data from neighboring countries.

In contrast, sub-lineage L1A exhibited a substantially more complex evolutionary history characterized by bidirectional trans-Pacific transmission. Consistent with previous studies, our analyses support a North American origin followed by introduction into East and Southeast Asia [7, 59]. Interestingly, Bayesian phylogeographic reconstruction revealed that the origin of sub-lineage L1A in Vietnam was the USA rather than China, despite their shared national borders. According to U.S. livestock international trade data, more than 16,500 hogs (including breeders) have been exported from the USA to Vietnam since 1996 (half of which were exported after the ASF outbreak), which could be the source of this introduction [60]. Interestingly, spatiotemporal analysis inferred a statistically supported migration event from Vietnam back to the USA around 2020. It was beyond the scope of this study to determine how this event may have occurred. Although this result should be interpreted cautiously due to incomplete sampling from intermediary countries, it nevertheless highlights the interconnected nature of global PRRSV evolution. International movement of feed ingredients (e.g., porcine endemic diarrhea RNA virus transmitted from Asia to the USA) [61], breeding genetics [59], germplasm [62], and complex multinational production networks may facilitate long-distance dissemination beyond direct live-animal trade. The Asia-to-U.S. transboundary spread event inferred by our analysis warrants further attention, as the USA looks to prevent the introduction of other viruses such as ASFV and foot-and-mouth disease virus.

From a disease control perspective, our findings reinforce the importance of integrating genomic surveillance into routine PRRS monitoring programs. Reliance on historical lineage classifications risks overlooking newly emerging virus groups that may differ in epidemiological behavior or vaccine responsiveness. Adoption of the standardized IPNC classification framework provides a consistent approach for identifying novel lineages and enables direct comparison among surveillance systems worldwide. Furthermore, because multiple emerging sub-lineages appear to have originated through repeated cross-border introductions, strengthening regional collaboration among Southeast Asian countries should become a priority. Improved sharing of sequence data, harmonized surveillance strategies, and coordinated monitoring of transboundary pig movement would substantially enhance early detection of emerging variants and improve preparedness for future disease incursions. The emergence and circulation of vaccine-like sequences identified in sub-lineages L5A and L8C also underscore the need to distinguish vaccine-derived viruses from field strains during molecular surveillance to avoid misinterpretation of epidemiological trends.

Several limitations should be acknowledged. First, all phylogenetic analyses were based on the ORF5 gene, which remains the international standard for PRRSV classification but represents only a small portion of the viral genome. Whole-genome sequencing would provide greater resolution for detecting recombination, identifying adaptive mutations, and resolving complex evolutionary relationships. Second, despite the implementation of a stratified downsampling framework designed to minimize sampling bias, publicly available sequence datasets remain unevenly distributed across countries and years, potentially influencing phylogeographic reconstruction and ancestral-state estimation. Finally, although Bayesian phylogeographic models provide valuable insight into historical viral movement, inferred transmission routes should not be interpreted as direct evidence of animal movement without complementary epidemiological and trade data.

## 5. Conclusions

The current study demonstrates that the ASF epidemic coincided with a profound restructuring of the Vietnamese PRRSV-2 population, characterized by persistence of the endemic sub-lineage L8E alongside repeated establishment of newly introduced sub-lineages. Prior to ASF, only sub-lineage L8E and vaccine-like sub-lineage L5A were present in Vietnam. In the post-ASF era, lineage diversity increased drastically, with six new (sub-)lineages present, resulting in at least eight (sub-)lineages co-circulating. The formal recognition of sub-lineages L1M and L10B expands the global PRRSV-2 nomenclature and highlights Vietnam as an important contributor to contemporary PRRSV evolution in Southeast Asia. Continued genomic surveillance, combined with standardized lineage classification and strengthened regional collaboration, will be essential to improve early detection of emerging variants and support evidence-based control of this continually evolving transboundary pathogen.

## Data Availability

The data that support the findings of this study are available from the corresponding author, Kimberly VanderWaal, upon reasonable request.

## Conflicts of Interest

The authors declared that there is no conflict of interest or personal relationship regarding the publication of this article.

## Author Contribution

T.C.N.: Conceptualization, Investigation, Formal analysis, Visualization, Writing – original draft manuscript; J.P.H.d.S and N.P.: Methodology, Data Validation, Writing – review and edit; R.T.: Supervision, Funding, Writing – review and edit; K.V.: Project administration, Conceptualization, Methodology, Data Validation, Writing – review and edit.

## Acknowledgements

The authors appreciate the Morrison Swine Health Monitoring Project (MSHMP) members. T.C.N. acknowledges the Second Century Fund (C2F) at Chulalongkorn University for his doctoral program scholarship and for the funding to conduct research abroad. This research is partially supported by the 90^th^ Anniversary of Chulalongkorn University Scholarship under the Ratchadapisek Somphot Endowment Fund and the National Research Council of Thailand: R. Thanawongnuwech NRCT Senior Scholar 2022 #N42A650553.

## Notes

### Competing Interest Statement

The authors have declared no competing interest.

