## Supplementary Files for "Shifts in Genetic Diversity of Porcine Reproductive and Respiratory Syndrome Virus 2 in Vietnam Before and After African Swine Fever: Increased Diversity and Novel Sub-lineages"

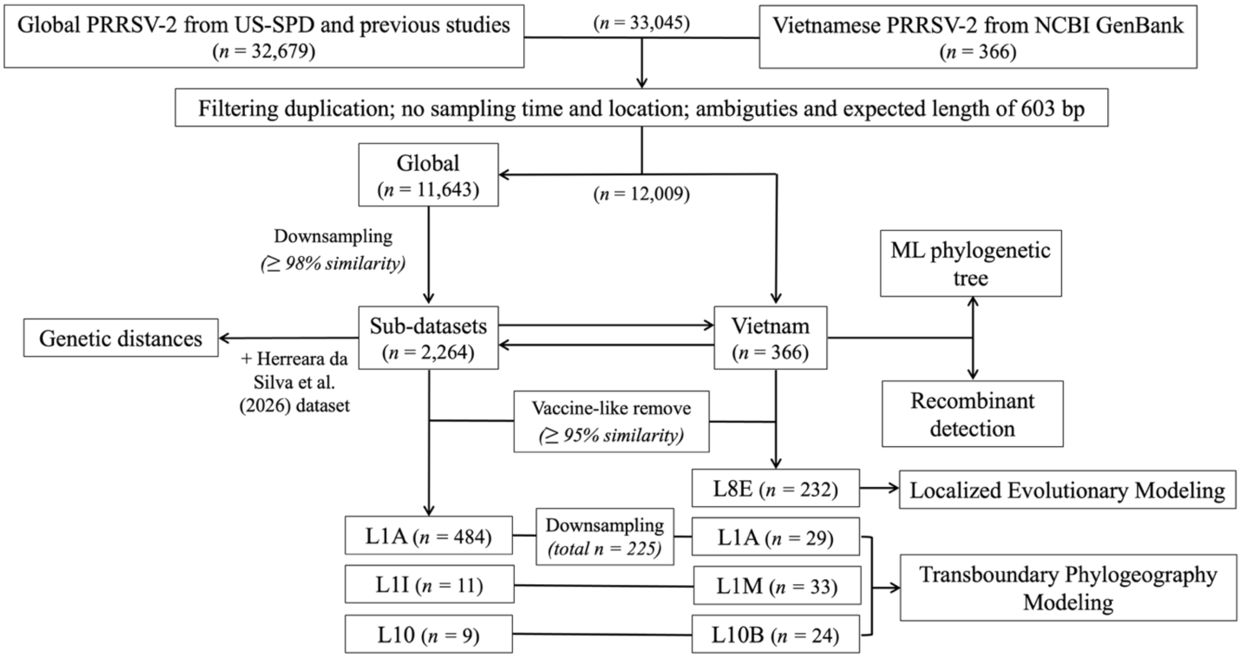


**Supplementary Figure S1.** Flowchart of the PRRSV-2 ORF5 data processing and analyses. Global (n = 32,679) and Vietnamese (n = 366) reference datasets were compiled and filtered to remove duplicates, unverified sampling metadata (time and location), ambiguous sequences, and fragments with an expected length of 603 bp. Global sequences underwent downsampling based on a genetic similarity threshold to construct representative sub-datasets. Maximum-likelihood (ML) phylogenetic trees were reconstructed, and recombination was detected from the combined global sub-datasets and the Vietnamese dataset. The major PRRSV-2 field sub-lineages in Vietnam were subjected to localized evolutionary modeling and transboundary phylogeography modeling. n: number of sequences.

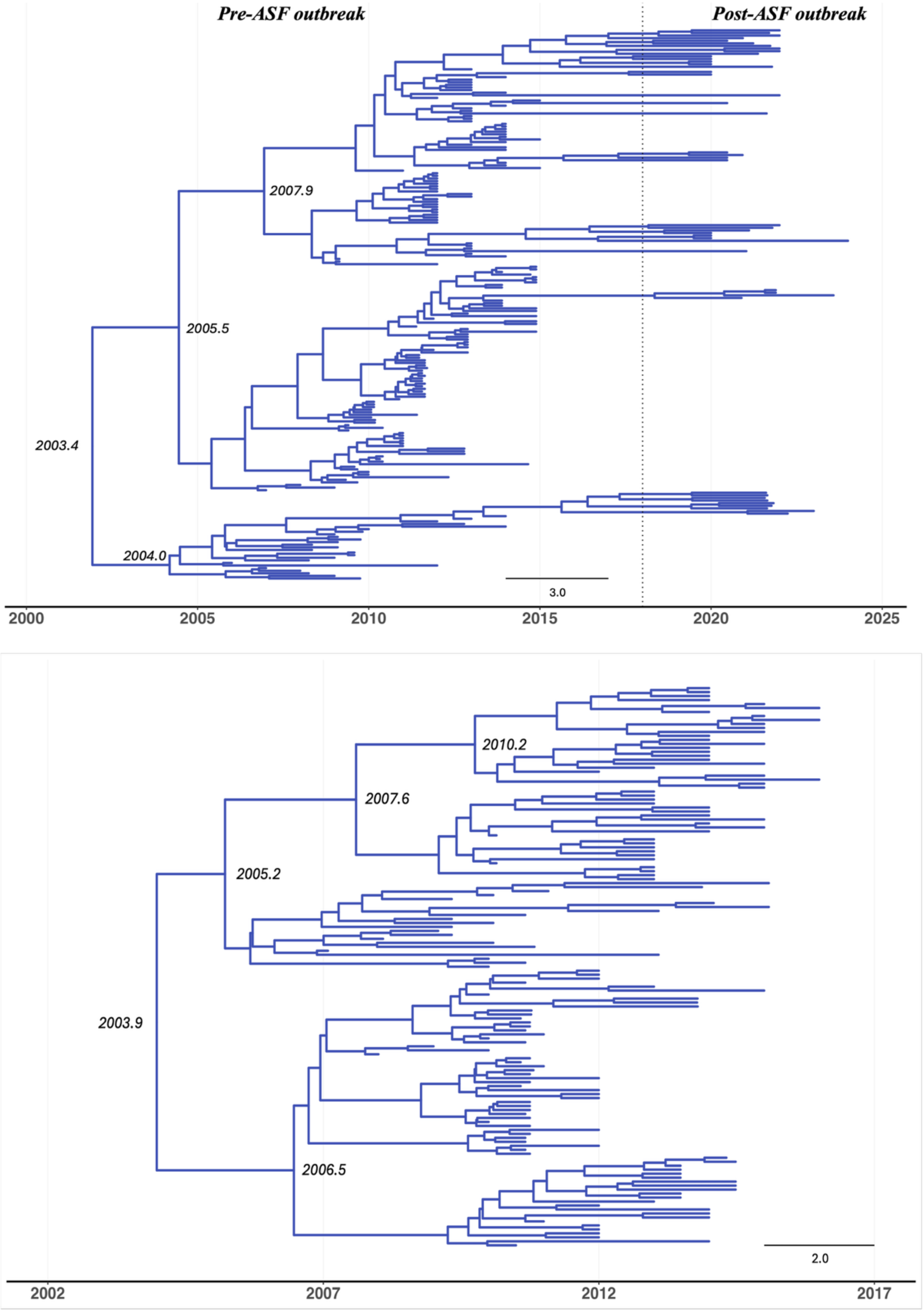


**(A)**

**(B)**

**Supplementary Figure S2.** Maximum Clade Credibility (MCC) tree of PRRSV-2 sub-lineage L8E in Vietnam **(A)** in the full longitudinal period (2007 – 2024) and **(B)** pre-ASF endemic (2007 – 2016). The tree was reconstructed using a Bayesian MCMC approach in BEAST v1.10.4. The dotted line represents the 2019 ASF outbreak in Vietnam. Internal node labels represent the estimated time of the Most Recent Common Ancestor (tMRCA) for major clades.

**Supplementary Table S1.** Table of genetic distances (Median [IQR: Q1 – Q3]) of two newly established sub-lineages and well-identified (sub-)lineages.

| (Sub-)lineage | L10B (L10-UD) | L1M (L1-UD) |
|---|---|---|
| **L1A** | 15.6 (14.9 – 16.3) | 14.2 (12.7 – 14.9) |
| **L1B** | 16.5 (15.9 – 17.1) | 14.8 (13.6 – 15.8) |
| **L1C** | 16.9 (16.2 – 17.4) | 14.9 (14.1 – 15.7) |
| **L1D** | 16.4 (15.6 – 17.1) | 14.9 (13.6 – 15.4) |
| **L1E** | 15.9 (14.9 – 16.9) | 15.4 (14.5 – 16.4) |
| **L1F** | 15.6 (15.1 – 16.2) | 14.7 (13.3 – 15.4) |
| **L1H** | 16.9 (16.2 – 17.7) | 15.0 (14.1 – 15.7) |
| **L1I** | 16.3 (15.5 – 17.0) | 13.9 (13.1 – 14.4) |
| **L1J** | 16.6 (15.8 – 17.2) | 15.1 (14.1 – 16.0) |
| **L1K** | 17.2 (16.4 – 17.9) | 15.7 (14.7 – 16.5) |
| **L1L** | 17.7 (17.0 – 18.3) | 15.8 (14.7 – 16.7) |
| **L1M (L1-UD)** | 18.4 (17.2 – 19.1) | 10.7 (7.3 – 12.2) |
| **L2** | 16.6 (15.8 – 17.3) | 15.6 (14.6 – 16.5) |
| **L3** | 17.2 (16.3 – 18.0) | 17.1 (16.2 – 17.7) |
| **L4** | 14.5 (13.6 – 15.4) | 14.6 (13.6 – 15.6) |
| **L5A** | 15.3 (14.4 – 15.9) | 15.4 (14.5 – 16.1) |
| **L5B** | 15.2 (14.7 – 15.7) | 16.6 (15.2 – 17.2) |
| **L6** | 16.1 (15.4 – 16.9) | 17.0 (15.9 – 17.7) |
| **L7** | 15.0 (13.4 – 15.7) | 15.4 (14.1 – 15.9) |
| **L8A** | 15.2 (14.6 – 15.8) | 15.2 (14.4 – 15.9) |
| **L8B** | 15.0 (14.2 – 15.7) | 15.6 (14.2 – 16.4) |
| **L8C** | 13.9 (13.4 – 14.5) | 14.4 (13.4 – 15.6) |
| **L8D** | 15.9 (15.3 – 16.4) | 17.6 (16.6 – 18.8) |
| **L8E** | 14.7 (14.1 – 15.1) | 15.7 (14.0 – 16.2) |
| **L9A** | 15.7 (14.8 – 16.7) | 15.6 (14.6 – 16.4) |
| **L9B** | 15.7 (14.9 – 16.6) | 16.4 (15.6 – 16.9) |
| **L9C** | 15.5 (14.9 – 16.2) | 16.3 (14.9 – 17.1) |
| **L9D** | 15.5 (14.7 – 16.6) | 15.8 (14.6 – 16.6) |
| **L9E** | 14.6 (13.7 – 15.4) | 15.5 (14.2 – 16.2) |
| **L10A (formerly L10)** | 12.7 (12.4 – 13.1) | 16.6 (14.4 – 16.9) |
| **L10B (L10-UD)** | 6.9 (4.0 – 10.0) | 18.4 (17.2 – 19.1) |
| **L11** | 16.3 (15.6 – 16.9) | 15.8 (15.0 – 16.6) |
| **L12** | 18.0 (17.2 – 19.0) | 17.0 (16.1 – 18.2) |
